# The spatial scale of adaptation in a native annual plant and its implications for responses to climate change

**DOI:** 10.1101/2022.01.22.477135

**Authors:** Amanda J. Gorton, John W. Benning, Peter Tiffin, David A. Moeller

**Author notes:** **Corresponding author:** John W. Benning. These authors contributed equally (Amanda J. Gorton, John W. Benning). **Author contributions** AJG, DAM and PT conceived the questions and experimental design. AJG performed experiments and contributed to data analyses. JWB conducted the data analyses in consultation with DAM and PT. AJG and JWB drafted the initial version of the manuscript. All authors contributed to interpreting data and editing the manuscript. **Data archiving:** Data and code are archived on FigShare: https://doi.org/10.6084/m9.figshare.c.6092868.v2.

## Abstract

Spatial patterns of adaptation provide important insights into agents of selection and expected responses of populations to climate change. Robust inference into the spatial scale of adaptation can be gained through reciprocal transplant experiments that combine multiple source populations and common gardens. Here we examine the spatial scale of local adaptation of the North American annual plant common ragweed, *Ambrosia artemisiifolia*, using data from four common gardens with 22 source populations sampled from across a ∼1200 km latitudinal gradient within the native range. We found evidence of local adaptation at the northernmost common garden, but maladaptation at the two southern gardens, where more southern source populations outperformed local populations. Overall, the spatial scale of adaptation was large — at the three gardens where distance between source populations and gardens explained variation in fitness, it took an average of 820 km for fitness to decline to 50% of its predicted maximum. Taken together, these results suggest that climate change has already caused maladaptation, especially across the southern portion of the range, and may result in northward range contraction over time.

## Introduction

Spatial variation in the selective environment can result in adaptive divergence among populations, i.e., local adaptation (Williams 1966). The extent to which populations are locally adapted depends on the balance between selection and gene flow, as well as exposure to genetic drift (Linhart and Grant 1996). Populations separated by short geographic distances are likely to experience high gene flow, which is expected to constrain local adaptation unless selection is very strong (Slatkin 1987; Lenormand 2002). However, nearby populations are also likely to share similar environments and experience more similar selection. Thus, nearby populations may not be expected to exhibit adaptive differences even in the absence of gene flow. As the distance between populations increases, gene flow is likely to decrease and environmental differences are expected to increase. Accordingly, the extent of adaptive differentiation and thus the signal of local adaptation may be expected to increase with geographical distance among populations (Slatkin 1985; Montalvo and Ellstrand 2000; Joshi et al. 2001; Hereford 2009). However, these expectations involve two assumptions: that geographic distance reflects ecologically relevant environmental distance, and that populations are (largely) in equilibrium with their environmental optima. These assumptions are challenged by some lines of empirical evidence, such as the presence of fine-scale (e.g., meter) adaptive differentiation (Antonovics and Bradshaw 1970; Janson 1983; Sambatti and Rice 2006), stochastic and directional temporal variation in environmental optima (e.g., Chevin et al. 2015; Bontrager et al. 2020; de Villemereuil et al. 2020), and the 30-50 percent of studies that find no evidence of local adaptation (Hereford 2009; Hargreaves et al. 2020; Runquist et al. 2020).

The most powerful method to identify patterns of adaptation is the common garden approach, where multiple source populations are planted into one site and performance is compared among populations. In such experiments, local adaptation manifests as higher fitness of “home” or “local” populations relative to foreign populations (Kawecki and Ebert 2004), and can be measured using categorical contrasts or, with enough populations, regressions of fitness on distance between a source population and garden (Fig. 1a). Many transplant experiments have been conducted, and in general, local adaptation seems to be common but certainly not ubiquitous (Hereford 2009; Runquist et al. 2020). However, there remain few studies that assess fitness differences among many source populations at multiple gardens along a broad environmental gradient (but see Colautti and Barrett 2013; Wilczek et al. 2014). Data from multiple common gardens across an environmental gradient can be used to evaluate the extent of local adaptation across a species’ range, as well as the strength of adaptation to large-scale selective agents like climate. Furthermore, sampling many populations allows for a quantitative, as opposed to qualitative, estimate of the spatial scale of adaptation (Fig. 1a), which is essential for understanding spatial patterns of adaptation (Galloway and Fenster 2000; Richardson et al. 2014). Though lifetime fitness is the best measure of adaptation in such studies, data on individual fitness components, such as early season survival or probability of flowering in plants, can provide insights into the life history stages underlying adaptation (Benning and Moeller 2021).

**Figure 1.**
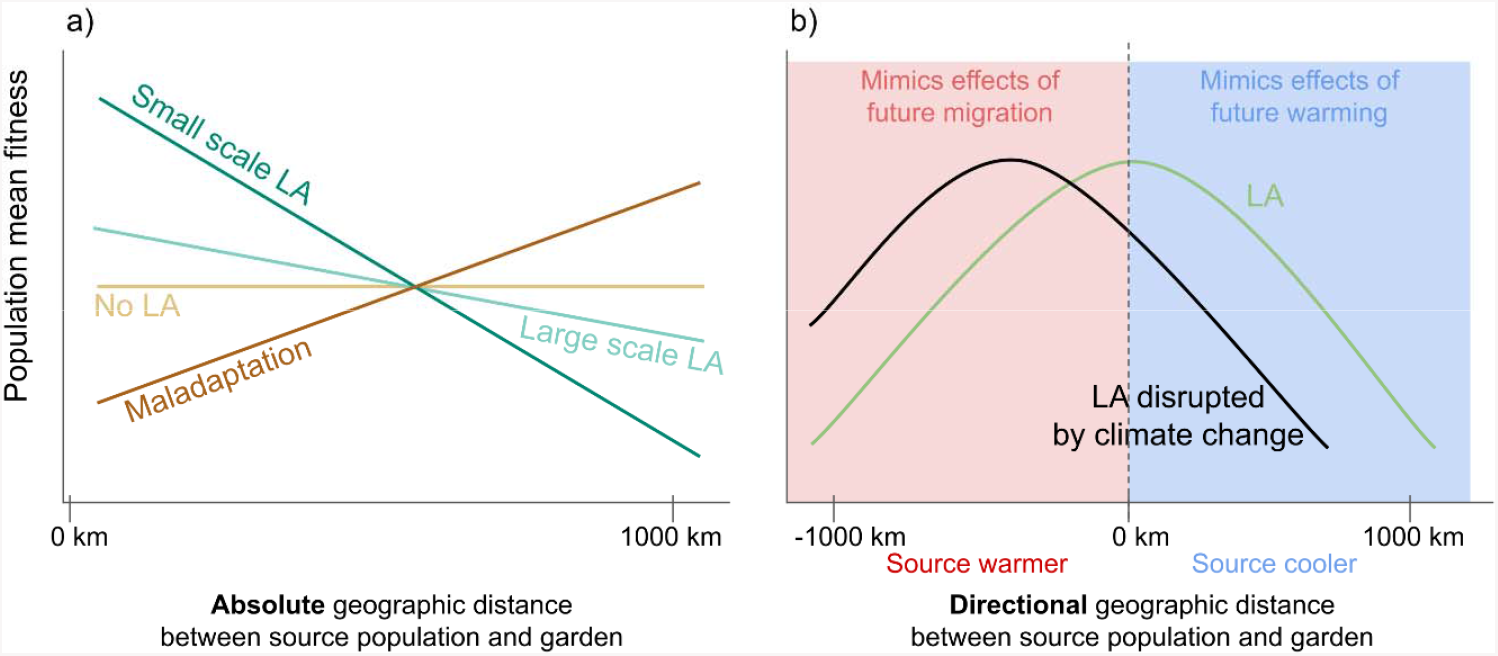
The scale and strength of adaptation can be estimated using regression analyses to characterize the relationship between the mean fitness of populations and geographic (or environmental) distances. **a)** The slope of the relationship reflects the geographic (or environmental) scale of adaptation: the more negative the slope, the more fitness is affected by a change in distance. A flat slope would indicate no local adaptation and a positive slope maladaptation (which could also vary in scale). The variance in fitness explained by the regression (R^2^) provides insight to the strength of local adaptation: a high R^2^ would indicate that the variable of interest is a strong predictor of fitness, and thus populations are strongly locally adapted. **b)** Shows how the relationship between fitness and a directional distance metric (such as latitudinal distance, calculated as *source population latitude - common garden latitude*) provides insight into whether local adaptation has been disrupted by climate change, and also offers predictions for population performance with future warming and migration. The green line is consistent with local adaptation at the location of an experimental garden (0 km). The black line, in which the populations with highest fitness were sourced from locations warmer than the garden, shows a pattern consistent with local adaptation having been recently disrupted by a warming climate. Populations transplanted into more poleward locations can be used to test for adaptation to sites potentially occupied after future migration, while transplanting populations into more equatorial gardens tests adaptation to warmer (future) temperatures.

In an effort to identify drivers of local adaptation, differences between source populations and common gardens are often characterized using geographic distance (Becker et al. 2006), temperature difference (Bontrager and Angert 2019), or multivariate environmental distance (Wright et al. 2017). Geographic distance is easy to measure and interpret and is often assumed to correlate with the amount of gene flow between populations (Slatkin 1987; Lenormand 2002), but is not a factor that drives local adaptation. Instead, because environmental variables are often spatially autocorrelated, geographic distance integrates multiple (known and unknown) environmental variables that drive adaptation. For instance, across large latitudinal scales geographic distance will capture variation in climate and photoperiod, both strong determinants of fitness in some species (Lacey 1988; Li et al. 2003; Fournier-Level et al. 2011; Keller et al. 2011; O’Brien et al. 2017; Campbell-Staton et al. 2018). Alternatively, distance can be calculated directly from environmental data, such as mean annual temperature or precipitation. Regardless of the variable used, distance between populations and gardens is usually calculated as absolute, Euclidean distance, and therefore does not differentiate between directional differences that may be important (but see Anderson and Wadgymar 2019). For example, transplanting a population 200 km poleward to a 2° C cooler location will likely have different effects on fitness than transplanting that same population 200 km equatorward to a 2° C warmer location. Incorporating such directional distances will be especially important for investigating patterns of adaptation when populations are potentially not in equilibrium with their environment due to perturbations like climate change (Fig. 1b).

Climate change is disrupting local adaptation due to shifting environmental optima (Anderson and Wadgymar 2019; Bontrager et al. 2020) and has already resulted in range shifts for some species (Chen et al. 2011; Lenoir et al. 2020). Experiments that assess fitness of populations sampled from across a climate gradient like latitude can provide two valuable insights: first, they can be used to estimate the strength of local adaptation – how much variance in fitness is explained by the distance between a population and garden? Second, with enough source populations, these experiments can quantify the spatial scale of local adaptation – what is the slope of the relationship between fitness and distance (Fig. 1a)? Across gradients where geographic and climatic distance strongly covary, a shallow relationship between fitness and geographic distance would suggest a large scale of adaptation to climate, such that species may be relatively unaffected by climate change. By contrast, a steep relationship between fitness and geographic distance would suggest that climate is an important driver of adaptation and populations are thus likely to be strongly affected by changing climate (Fig. 1a). When analyzed with directional measures of distance, such experiments can also uncover evidence that climate warming has already disrupted local adaptation, for instance if populations from warmer regions have higher fitness than local populations (Gorton et al. 2019; Bontrager et al. 2020; Fig. 1b).

Furthermore, if populations are sourced from regions colder than the common garden, the transplant experiment can be thought of as a space-for-time substitution with respect to climate change (e.g., Etterson and Shaw 2001; Fig. 1b). By evaluating the performance of “cold” populations in a warmer locale, one can predict fitness of these populations in their home sites with future warming (though this design does not account for other environmental factors differing between locales). Moreover, populations sourced from regions warmer than the common garden can mimic potential future migration as populations track shifting climatic isotherms. In this case, fitness in the common garden would reflect adaptation to potentially important non-climatic aspects of a future environment that will likely not shift with climate change, such as day length or soil type.

We used a reciprocal transplant experiment of a North American annual plant, *Ambrosia artemisiifolia* (common ragweed), to quantify the strength and spatial scale of local adaptation and explore the consequences of a warming climate across a large portion of the species’ geographic range. The experiment included four common gardens, in which we planted individuals from 22 populations collected from across a ∼1200 km latitudinal gradient. Our main questions were:

1. To what extent are populations locally adapted or maladapted across the species’ range?
2. What distance metric [geographic (absolute or directional), climatic (absolute or directional)] is the strongest predictor of fitness variation among populations?
3. What is the spatial scale of adaptation to climate?
4. Which life history stages underlie adaptation and how do these vary across regions of the species’ range?

## Methods

### Study system

*Ambrosia artemisiifolia* (common ragweed; Asteraceae) is a self-incompatible (Friedman and Barrett 2008), monoecious, wind-pollinated annual plant (Jones 1936; Essl et al. 2015). The species is found throughout North America, excluding the northernmost portion of Canada, and is invasive on several continents (Genton et al. 2005; Chauvel et al. 2006; Xu et al. 2006). Within North America the species’ abundance increased dramatically as agriculture spread across the continent (Grimm 2001; Martin et al. 2014) and today common ragweed is most abundant in the eastern and midwestern United States and Canada (Kartesz 2013). It is widely distributed across broad climatic gradients, making it a useful system to investigate questions regarding the scale of adaptation to climate. Latitudinal clines in flowering time have been documented on multiple continents, with plants from higher latitudes flowering earlier (measured by Julian date) than plants from lower latitudes in both common gardens and in the field (Hodgins and Rieseberg 2011; Leiblein-Wild and Tackenberg 2014; Gorton et al. 2019). These clines suggest that populations are adapted to local climate, even though transcriptome data revealed that population structure in North American ragweed only occurs at broad spatial scales (Hämälä et al. 2020). Phenotypic differentiation among ragweed plants sampled from 1 – 3 km distances (Kostanecki et al. 2021) suggests that adaptive divergence likely also occurs at finer spatial scales.

*A. artemisiifolia* is a ruderal plant that is often abundant in open, disturbed habitats such as riverbanks, roadsides, edges of agricultural fields, and urban areas. It is a summer annual that typically germinates in late spring (May – June) and flowers in late summer or early fall (August – Sept). The transition to reproduction is cued by shortening day lengths after the summer solstice, and the onset of frosts in the fall terminates the growing season. Staminate capitula (*i*.*e*. male flowering heads) are found in racemes, which produce pollen that is one of the primary causes of summer and fall allergic rhinitis. Pistillate capitula (*i*.*e*. female flowering heads) are found in axillary clusters below the male flowers; each individual flower develops into an achene (a small, single seeded fruit) that readily falls off the plant once ripe. These axillary clusters contain on average between 1-5 achenes or seeds (Essl et al. 2015). Hereafter we refer to these groups of achenes as ‘fruits’; thus, a fruit count of one corresponds to between one and five seeds.

### Seed collections

During October 2015 – January 2016, we collected seeds from 22 populations of *A. artemisiifolia* across a region spanning 11° latitude (∼1200 km) and 7° longitude (∼550 km) (Table S1, Figure 2). We prioritized sampling across a replicated, broad latitudinal gradient in order to directly address questions concerning effects of climate warming as outlined in the Introduction and Figure 1b (note that because our study occurred in North America, when we use “southern” and “northern” below, this refers to equatorward and poleward locations, respectively). Sampling locations were an average of 290 km apart along two latitudinal transects. At most latitudes, along each transect, we sampled two populations separated by 50 – 60 km. The populations were collected from a range of habitats, including abandoned lots in urban areas, roadsides, river edges, and parks. At each sampling location we collected seeds from 16-25 maternal plants, each separated by at least 3 m.

**Figure 2.**
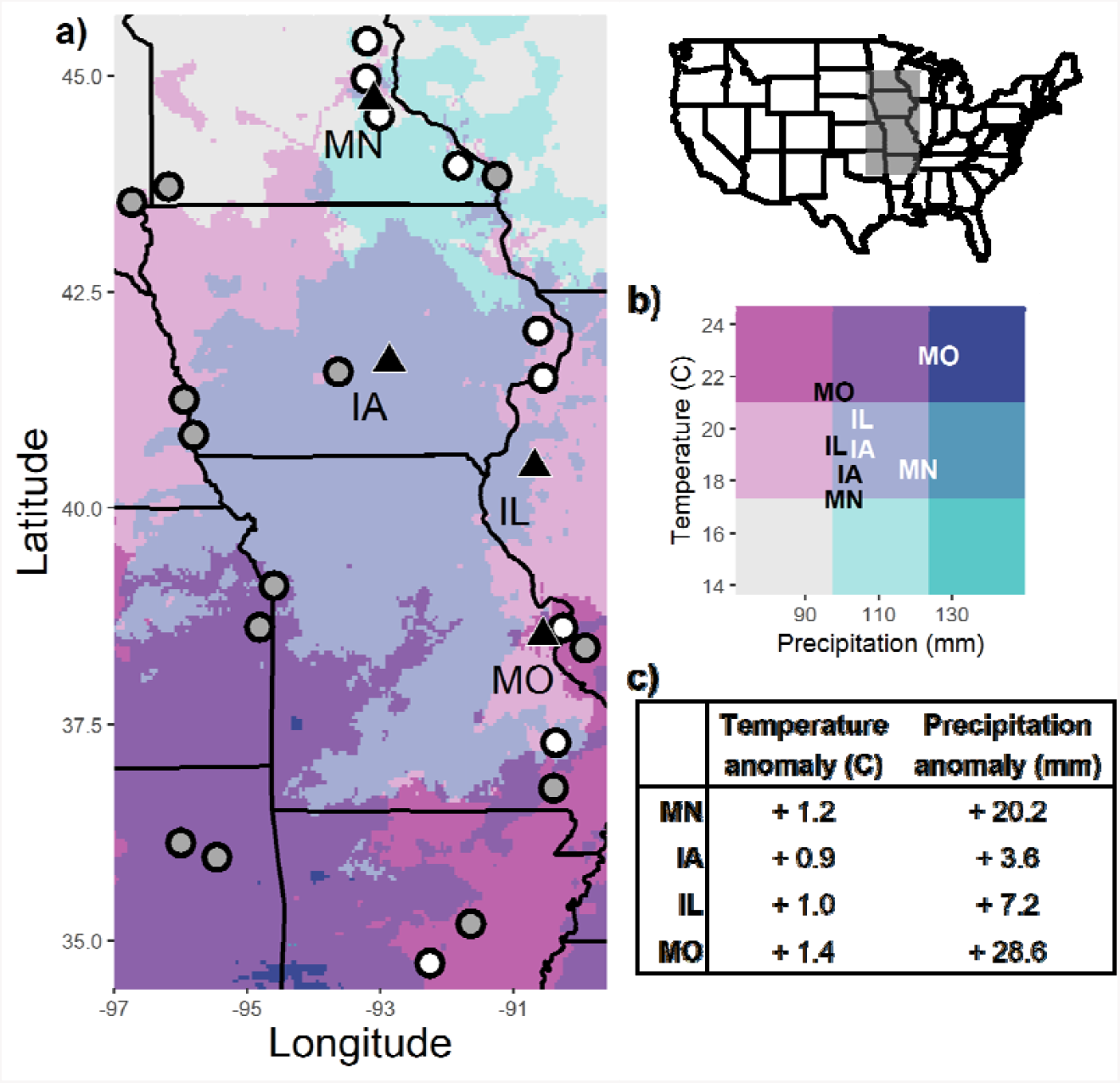
a) Map showing the locations of the 22 source populations (circles) and four common gardens (triangles). White circles show the nine focal populations for which complete lifetime fitness data were collected; grey circles show the remaining 13 populations where final fruit counts were not obtained. b) Climatic regions based on temperature and precipitation representative of the growing season (May - October), averaged over years 1986-2016. Climate data were extracted from PRISM. Garden names are plotted on the color legend according to their historical climate averages (1986-2015, black text) and experiment year weather (2016, white text). c) Table shows how temperature and precipitation during the experiment year (2016) differed from the long-term average (1986-2015).

### Reciprocal transplant experiment

In Spring 2016, we planted four common gardens (hereafter ‘gardens’) that were separated by an average of 260 km (range: 218 – 338 km) and differ in annual temperature, precipitation, and photoperiod (Figure 1). The gardens were in the states of Minnesota (MN), Iowa (IA), Illinois (IL), and Missouri (MO) (see Table S2 for exact locations). The soil in all gardens was a silty loam.

Between May 15 and May 30, 2016, we planted the four gardens, starting at the southernmost site (MO) and ending in the northernmost site (MN). The timing of planting approximately matched the germination timing of natural populations at each latitude (Gorton, personal observations). Prior to planting, each garden was sprayed with glyphosate and tilled, creating an environment similar to the disturbed, low-competition habitats where *A. artemisiifolia* is commonly found. *A. artemisiifolia* has the potential for long-term dormancy, with seeds in the seed bank remaining viable for many years (Toole and Brown 1946). To minimize confusion between experimental plants and those that emerged from the seed bank, we used a ProPlugger to remove a soil plug (5 cm x 5.5 cm) at each planting spot. The holes were then filled with Berger germination mix (Berger, Quebec).

Prior to planting, all seeds were stratified in moist silica sand in the dark at 4°C for 10 weeks to break dormancy (Willemsen 1975). The seeds were planted in a randomized complete block design (*n* = 25 blocks), so that each block contained seeds from each population (22 populations × 25 plants per population = 550 plants/ garden). For each source population, we pooled an equal number of seeds from 12 – 25 maternal plants (Table S1). At each planting spot, we planted 2 – 4 stratified seeds directly into the ground and watered the seeds immediately after planting. Each garden had 55 rows and 10 columns of plants, planting spots were spaced 15 cm apart along each row, and the spacing between columns alternated between 30 and 80 cm. Four weeks after planting, plants were thinned to a single seedling per planting spot. Throughout the growing season, all gardens were regularly weeded to reduce competition from non-experimental plants.

### Fitness components

We visited each garden 4, 8, 12, and 16 weeks after planting, and at the end of the growing season. During each week, we documented survivorship and height, and scored each plant for developmental stage (e.g. vegetative, flowering, fruiting, etc.), male flowering stage, female flowering stage. For each of the three later traits, each plant received a score between one and five, corresponding to different reproductive or flowering stages (scoring categories presented in Table S3); plants could receive intermediate scores (e.g., 3.5). We estimated female fitness at each garden by counting the number of mature fruits on 12-15 randomly selected plants from each of 9 of the 22 source populations (108-135 plants per garden) at 16 weeks and at the end of the season. These nine populations span the full latitudinal range of the source populations (Fig. 2, Table S1). Due to the distance between gardens, high fecundity, and the narrow time window in which data needed to be collected, it was not possible to quantify female fitness for every plant. For those plants for which we quantified female fitness, we counted the number of fruits on every fourth branch (ca. 20 – 50 plant^-1^), starting with the lowest branch, and moving up towards the apical meristem. We subsampled branches in this way to account for variation in allocation to female reproduction across plant development. We multiplied this number by four to get a whole-plant estimate of female fitness. We also recorded fruit maturity on each group of fruits. Fruits were considered mature and given a score of one if at least one achene in the axillary cluster was brown. Plants with only immature green fruit were given a fitness score of zero, as these fruits were unlikely to mature prior to the first frost. Hereafter ‘number of fruits’ refers only to mature fruits. We collected data on female fitness (number of mature fruits produced) at both 16 weeks after planting and at the end of the growing season to capture the range in life span of the different populations, day length, and the length of the growing season across gardens. This was particularly relevant at the southern gardens, IL and MO, which had longer growing seasons. We counted fruits on the same individuals at both visits, and where possible, the same branches. When fruit counts differed across visits, we kept the higher value. It was possible for an individual to have a higher fruit count at the first census because some individuals had senesced already due to a shorter lifespan by the second census, leading to the mature fruits dropping off the plant.

We used life history data (developmental stage at 12 weeks, female flowering stage at 12 weeks, height at 16 weeks, fruit number) collected on plants from the 9 focal populations (N = 524) to build predictive models of individual fitness based on phenotypic data (SI: Predicting fitness). These models accounted for a large proportion of the variation in observed population mean fitness, with a correlation of *r* = 0.94 between predicted and observed population mean fitness (SI: Predicting fitness). We used these models to predict fitness of the individuals from the remaining 13 populations (and the individuals from the 9 focal populations for which we did not have final fitness data). Results from regressions of *fitness ∼ distance* using data from either the 9 or all 22 populations were very similar (SI: Predicting fitness; Fig. S3); we present results using the 22 populations because these capture a wider geographic sampling of populations and increase statistical power. Thus, the final dataset consisted of population mean fitnesses calculated from 1) observed individual fitness data for individuals from the nine focal populations for which we recorded fruit counts in the field, and individuals that died, and 2) predicted individual fitness for the remaining individuals where final fruit counts were not recorded. We do note that predicting fitness in this way assumes that the relationship between phenotypes and fitness in the nine focal populations holds for the other 13 populations (SI: Predicting fitness; Fig. S2).

### Statistical analyses

To test for local adaptation at each garden (question 1 from Introduction), we used linear regression models to estimate the relationship between mean population fitness (measured as mean individual fruit count) and Euclidean geographic distance (between source population and garden). We used analogous regression models to compare the explanatory power of different distance predictors (absolute geographic, directional geographic, absolute climatic, and directional climatic) for fitness (question 2). We calculated the spatial scale of adaptation to climate (question 3) using the estimated slopes from these models. To identify specific life history stages that showed the strongest signal of divergence among populations (question 4), we used generalized linear and linear models to estimate the relationship between separate fitness components (as opposed to lifetime fitness) and distance. The unit of analysis in all models below was population mean fitness (i.e., mean number of fruits produced by individuals). Analyses were conducted in R (v. 4.1.2, R Foundation For Statistical Computing 2021) and all data and code to replicate the analyses are available at FigShare (https://doi.org/10.6084/m9.figshare.c.6092868.v2). We used tidyverse packages (Wickham et al. 2019) for data manipulation and visualization, along with visreg (Breheny and Burchett 2017) for model visualization.

### Distance between populations and gardens

In common garden experiments such as ours, local adaptation should manifest as a negative relationship between population mean fitness and the geographic distance between source population and the common garden (Fig. 1a). Geographic distance is most often measured as the Euclidean distance between population and garden, which is the method we use for testing local adaptation at each garden (*Evidence for local adaptation across gardens* below). This distance measure, however, fails to account for directionality that may be important in explaining fitness variation among populations — e.g., the environment a fixed distance north of a garden is likely to be quite different from the environment that same distance south of a garden. For this reason, we also calculated directional geographic distance measures, which were the differences in latitude and longitude between source populations and gardens. To make Euclidean distance more directly comparable to directional geographic distance, we decomposed Euclidean distance into the absolute values of latitudinal and longitudinal distances between populations and gardens. Hereafter, geographic distance refers to the combination of latitudinal and longitudinal distance either for absolute or directional measures. To aid in interpretability, we transformed these distances in latitude/longitude to distances in kilometers for visualization.

Environmental distance is an alternative to geographic distance. Because we were interested in assessing adaptation to climate, we calculated environmental distance between each garden and source population as climatic distance using monthly mean temperature and monthly total precipitation data from 1986-2016 extracted from PRISM (Oregon State University 2020). We extracted monthly climate data for the growing months (May – Oct) for each source population from 1986 – 2015, and for each garden in 2016 (the experiment year). We then calculated mean monthly temperature and precipitation for each year as the average across the six growing months. For source populations, mean temperature and precipitation were averaged over years 1986 – 2015. We then calculated temperature distances as the difference between a source population location’s long-term mean temperature and the experiment year (2016) temperature of the common garden. Precipitation difference was calculated in the same fashion. We calculated both absolute and directional climatic distances in the same way we calculated geographic distances. See Table S4 for correlation matrix of all distance metrics.

### Evidence for local adaptation across gardens

We used linear regression to regress population mean fitness (mean fruit set of all individuals in a population) on Euclidean distance between populations and gardens (e.g., *lm(fitness ∼ Euclidian distance)*. Local adaptation would be indicated by a negative relationship between fitness and distance (Fig. 1a).

### Comparing distance predictors of fitness variation

To compare the explanatory power of Euclidean distance against different population-garden distances (absolute geographic, directional geographic, absolute climatic, and directional climatic), we used the same linear regression framework as above. Specifically, we used linear regression to regress population mean fitness on the four sets of distance metrics separately (e.g., *lm(fitness ∼ directional latitudinal distance + directional longitudinal distance)*. We then estimated the proportion of among-population variance explained by each of the distance metrics using ANOVA with Type II SS [Anova() function in the car package (Fox and Weisberg 2019)]. Because the *a priori* expectation for the relationship between fitness and our directional distance metrics (directional geographic and climatic) is quadratic (Fig. 1b), we included quadratic terms in these models.

To select parsimonious final models for downstream analyses (below), we removed terms if ANOVA indicated they did not explain significant variance in fitness (*P* > 0.05). We then chose the distance model with the highest explanatory power (highest average *R*^2^ across gardens) to evaluate the scale of adaptation in the four gardens and test the influence of different life history stages on patterns of adaptation.

### Scale of adaptation

We quantified the scale of adaptation at each garden as the slope of the linear regression of fitness on distance, using the distance metric that explained the most variation in fitness (see above). This slope thus represents the predicted change in fitness with distance at each common garden. To test whether the scale of adaptation differed among gardens, we built an omnibus linear regression model (*lm(fitness ∼ garden × distance)*). To account for fitness variation among gardens, we mean-relativized population fitnesses within gardens. We used ANOVA with Type II SS to test whether slopes differed among gardens. Prior to analyses, we transformed distances for the MN garden into their additive inverses (i.e., multiplied by -1) so that the sign of the slope was the same across all gardens (see Results). This allowed us to test whether the absolute value of slopes differed among gardens (as slopes of e.g., *m* = −5 and *m* = 5 would indicate the same spatial scale of adaptation). Pairwise contrasts of slopes between gardens were performed with the emtrends function in the emmeans package (Lenth 2018), with a Tukey adjustment for *P* values.

### Conditional fitness component analyses

The above analyses identify the relative ability of different distance metrics to explain lifetime fitness variation, but they do not identify the life history stages that underlie adaptation. To identify those stages, we analyzed fitness components (probability of survival to 4 weeks, probability of fruiting, and number of fruits) to identify which life history stages most strongly drive overall lifetime fitness. For the subset of plants for which fruits were not counted, we used the predicted values from phenotypic models (as in lifetime fitness analyses, SI: Predicting fitness). For probability of mature fruits, all individuals scored as having at least one mature fruit were assigned a value of 1 and others were assigned a value of 0. Analyses were conditional such that individuals were included in the analysis only if they reached the previous life history stage (i.e., only plants that survived to 4 weeks were included in the analyses for probability of fruiting; number of fruits were modeled only for plants that produced fruits). We used logistic regression on population-level data (*i*.*e*., each data point was number of successes and failures per population) to estimate the influence of the distance metric that explained the most variation in lifetime fitness regressions on probability of early survival and probability of fruiting; *glm(cbind(successes, failures) ∼ distance, family = binomial)*. We used linear regression on population means to estimate the influence of the same distance metric on the number of fruits; *lm(number of fruits ∼ distance)*.

## Results

### Fitness and local adaptation across gardens

Growing season temperature and precipitation were higher than the historical average at all gardens (Fig. 2). Ragweed fitness differed starkly among gardens, with the lowest fitness (mean of 173 fruits per plant) at the northernmost garden, MN, and highest fitness (968 fruits per plant) at IL, the garden near the center of the sampled populations (Fig. 3). There was considerable among-population variation in fitness at each of the four gardens. However, we found evidence for local adaptation only in the northernmost (MN) and southernmost (MO) sites, where there were negative relationships between fitness and Euclidian distance between source populations and gardens (Fig. 3a, Table 1). The strength of these relationships was only moderately strong (*R*^2^ of 0.20 at MN and 0.27 at MO), indicating considerable among-population variation in fitness that was not related to Euclidean distance. Non-local populations often had higher fitness than local populations, especially at MO. At IA and IL, there was little relationship between fitness and Euclidean distance from the garden (Fig. 3a).

**Table 1.**
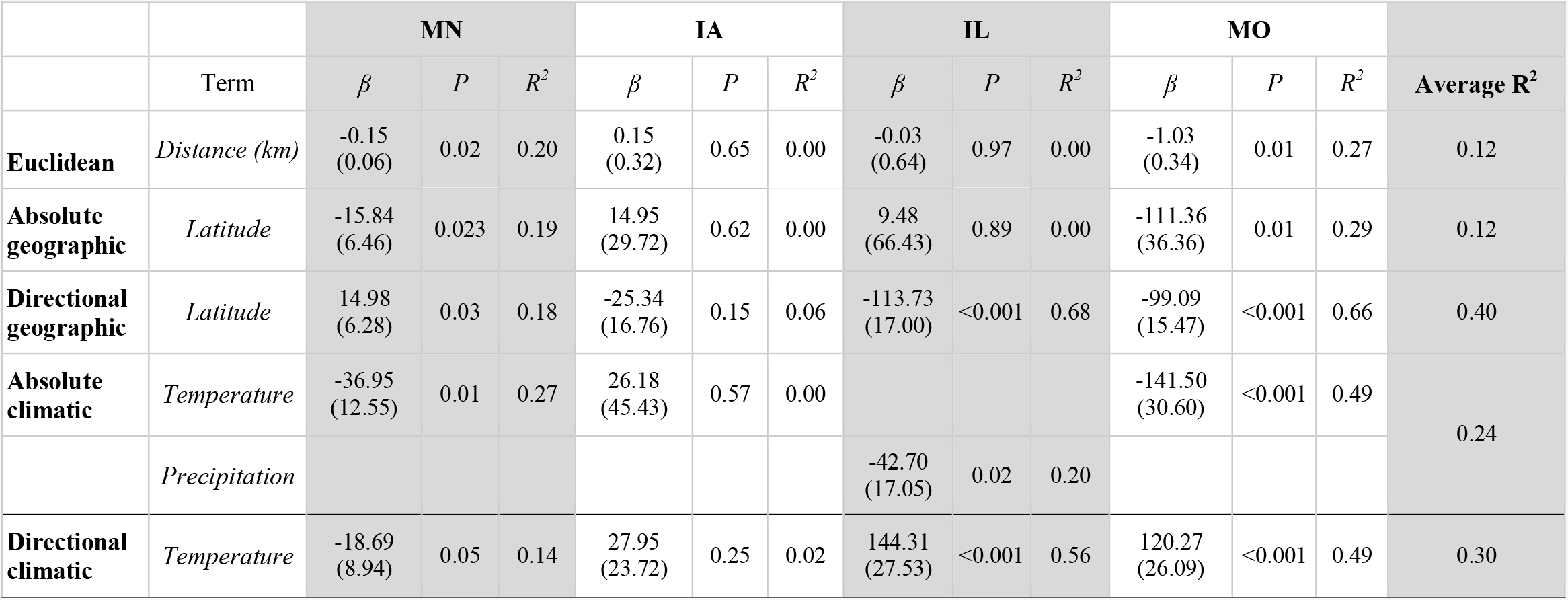
ANOVA tables for final models of the linear regressions of *fitness ∼ distance* at each garden. Reported are adjusted *R*^2^. All *P* values are m from F tests with 1, 20 df. ANOVA tables for full models including all predictors are in Table S5.

**Figure 3.**
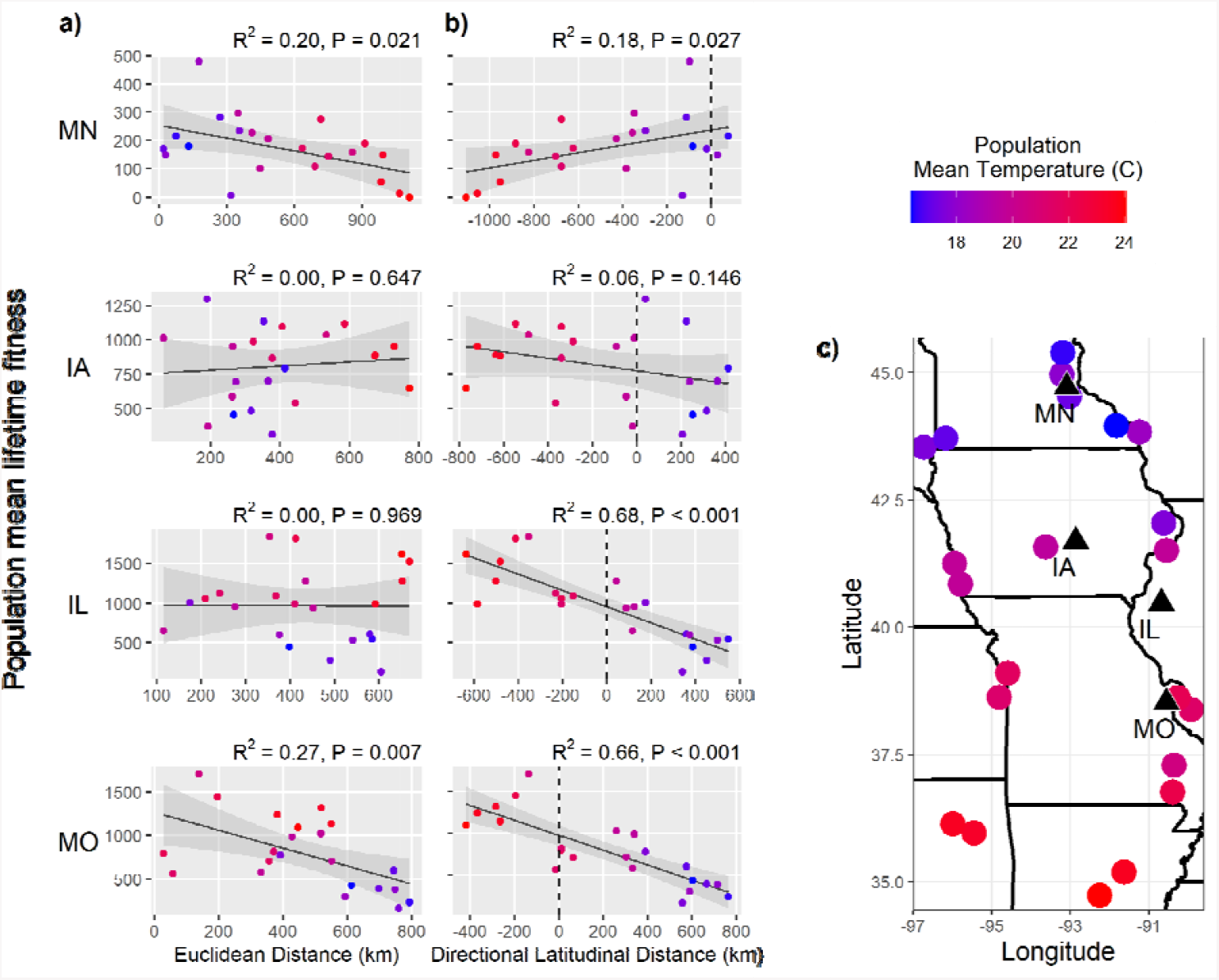
Linear regressions of mean lifetime fitness (number of fruits) on **a)** Euclidean distance and **b)** directional latitudinal distance between source populations and gardens. Negative directional latitudinal distance indicates a population sourced from a latitude south of the garden. Shaded ribbon in all plots is the 95% confidence band; dotted line in (**b**) represents garden location (i.e., is at 0 km distance). Each point represents mean fitness of a single population and is colored by population growing season mean temperature (averaged 1986-2015). Latitudinal distance was transformed to distance in kilometers based on 1° ≈ 111 km. Garden and population locations **(c)**, with populations colored by population growing season mean temperature (averaged 1986-2015).

### Predictors of fitness

We compared the explanatory power of absolute geographic, directional geographic, absolute climatic, and directional climatic distances for explaining fitness variation among populations at each common garden. Both geographic distance metrics comprised longitudinal and latitudinal distance, but because longitudinal distance contributed little explanatory power (Table S5), we focus on regressions of *fitness ∼ latitudinal distance*. Absolute and directional precipitation distances contributed little explanatory power to our climatic models (Table S5). Directional temperature distance explained substantial variation in fitness at the gardens, but because directional temperature and directional latitudinal distance were tightly correlated (*r* = -0.95; Table S4), we focus on latitudinal distances below but include temperature information in figures and text to aid in interpretation (full climatic distance results can be found in Tables 1 and S5). Quadratic distance terms used in directional distance models contributed little explanatory power (Table S5).

Averaged across gardens, directional latitudinal distance was the best predictor of population mean fitness (mean R^2^ = 0.40), and greatly outperformed absolute measures of distance (Fig. 3; Tables 1, S5). However, the better predictive ability of directional latitudinal distance was primarily driven by the two southern gardens, IL and MO. Absolute and directional latitudinal distance explained similar amounts of fitness variation at MN (R^2^ = 0.19 and 0.18, respectively), which is expected given that the MN garden is near the northern edge of the population sampling area and absolute and directional latitudinal distance were almost perfectly correlated (*r* = -0.996, Table S4). Neither distance metric explained an appreciable proportion of fitness variation at IA (both R^2^ < 0.07, P > 0.1). At the two southernmost gardens, IL and MO, population mean fitness was much better predicted by directional than by absolute latitudinal distance. The importance of using directional distance was particularly apparent at IL – whereas absolute latitudinal distance explained minimal fitness variation among populations at IL (adjusted R^2^ = 0.00), directional latitudinal distance predicted fitness variation very well (R^2^ = 0.68). At both IL and MO, there was a strong trend of populations originating from warmer and more southern regions than the gardens having the highest mean fitness.

### Spatial scale of adaptation

We quantified the spatial scale of adaptation at each garden as the slope of the linear regression of population mean fitness on directional latitudinal distance (Fig. 3b). This slope thus represents the predicted change in fitness with distance at each common garden. Slopes were generally shallow, indicating a large spatial scale of adaptation (mean absolute slope averaged across gardens = 57 fruits 100 km^-1^; ca. 5 – 10% of mean total fruit production, depending on the garden). There was substantial variation in slopes among gardens when modeled using absolute fitness (slope ± SE: MN = -13.5 ± 6 fruits 100 km^-1^; IA = 22.8 ± 15; IL = 102.5 ± 15; MO = 89.3 ± 14). When we scaled garden mean fitness to account for fitness variation among gardens, absolute slopes did differ among gardens (*garden* × *directional latitudinal distance* F_3,80_ = 2.99, *P* = 0.036), but this was driven by the IA garden having a comparatively shallow slope. Slopes were statistically indistinguishable (all Tukey-adjusted *P* > 0.6 for pairwise contrasts) in the three gardens where directional latitudinal distance explained appreciable fitness variation among populations (absolute slope ± SE: MN = 0.09 ± 0.03; IL = 0.12 ± 0.03; MO = 0.13 ± 0.03).

### Life history stages underlying fitness variation

To identify the life history stages most strongly affecting lifetime fitness differences among populations, we partitioned lifetime fitness into three stages: probability of surviving to four weeks post-planting, probability of fruiting given that a plant survived, and number of seeds produced given that a plant fruited. We found that early plant survival was high for all populations and all gardens and thus contributed little to either among-population or among-garden differences in plant fitness (Fig. 4). Similarly, at the three most southern gardens, nearly all plants that survived fruited. By contrast, there was a strong relationship between probability of fruiting in the MN garden and directional latitudinal distance, where plants from lower latitudes generally had delayed phenology (Fig. S4) and were much less likely to produce mature fruits before the end of the growing season (Fig. 4). The most consistent contributor to among-population fitness differences was number of fruits produced (i.e., fecundity) – at MN, IA, and IL, populations sampled from more southern locations tended to produce more fruits per individual than populations sampled from more northern locations (Fig. 4).

**Figure 4.**
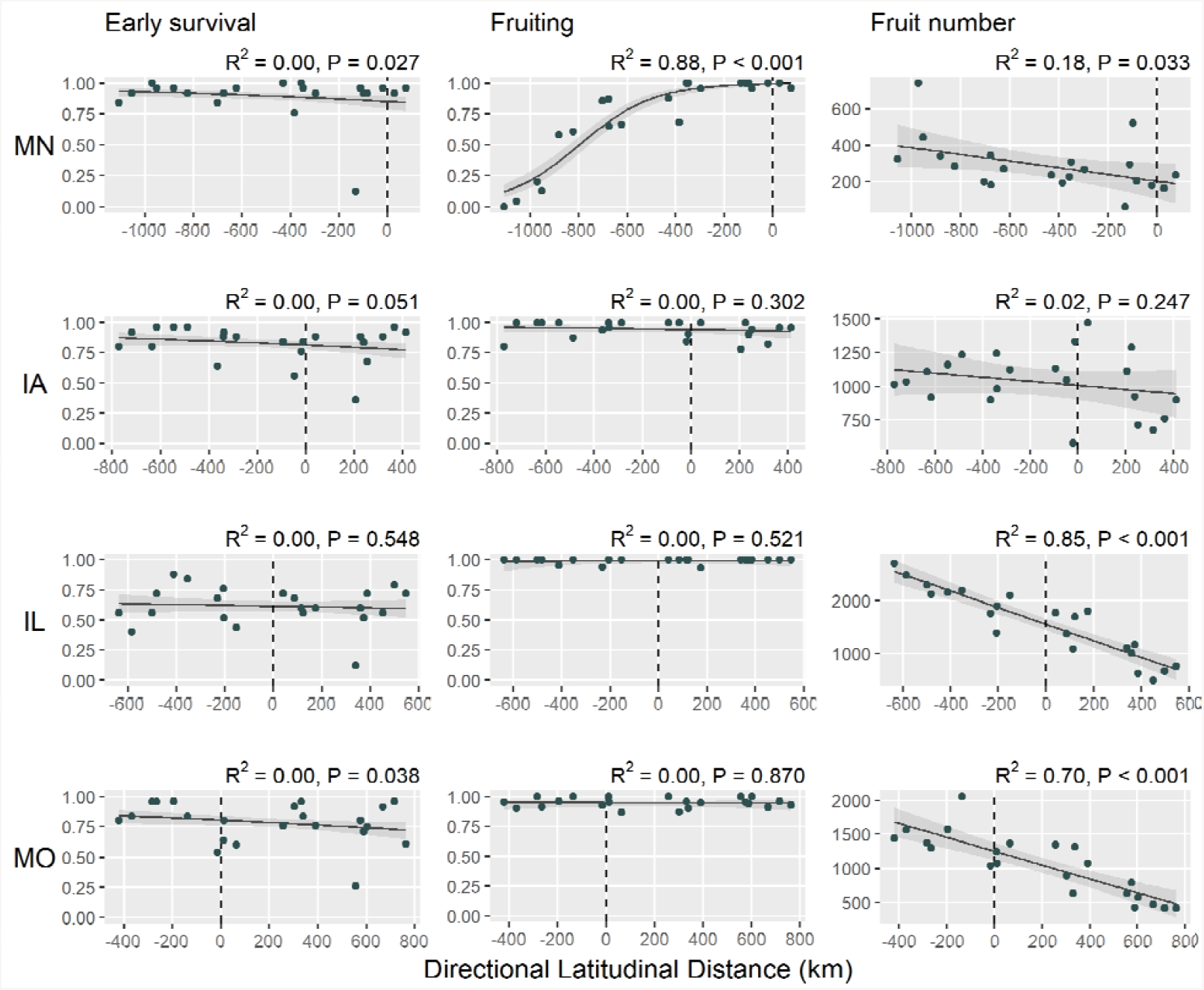
Relationships between conditional fitness components [*P*(early survival), *P*(fruiting), and number of fruits] and directional latitudinal distance for each common garden. Early surviva and fruiting were modeled with logistic regression, fruit number with linear regression. ANOVA results are in Table S6. Points represent population means. Shaded ribbon is the 95% confidence band, and the dotted line represents garden location (i.e., is at 0 km distance). For logistic regressions, the pseudo-R^2^ is based on differences between measured population proportions and model-predicted probabilities. Early survival was measured for all plants; fruiting and number of fruits were predicted from phenotypic models for the subset of plants where these traits were not measured (SI “Predicting fitness”). Latitudinal distance was transformed to distance in kilometers based on 1° ≈ 111 km.

## Discussion

The study of local adaptation provides insight into the spatial scale of adaptive differentiation, the influence of selection versus other processes on evolution, and the environmental drivers of population differentiation (Savolainen et al. 2007; Whitlock 2015; Runquist et al. 2020). Understanding geographic patterns of fitness and the scale of adaptation is also central to predicting how populations will respond to a changing climate (Aitken et al. 2008; Anderson 2015). To understand spatial patterns of adaptation to climate in common ragweed, we performed a reciprocal transplant experiment at four common gardens with 22 populations sampled across a large portion of ragweed’s geographic range in the central United States. We found that: (1) there was evidence for local adaptation at the northern edge of the range, but not at the other gardens; (2) directional latitudinal distance between populations and gardens was the best predictor of population mean fitness; (3) the scale of adaptation to climate in ragweed is large, but climate change has already caused some populations to be in disequilibrium with their environment, with southern populations outperforming local populations; (4) the life history stages underlying adaptation were strongly dependent on the location of the common garden within the species’ range. Our results suggest that climate change has already disrupted local adaptation in some regions and emphasize that directional measures of distance may often be essential for understanding patterns of fitness and predicting species’ responses to climate change.

The spatial scale of adaptation reflects a balance between local selection pressures driving adaptive divergence and gene flow homogenizing populations (Slatkin 1987; Richardson et al. 2014). Although many transplant experiments have been conducted, only a few have provided a quantitative assessment of the spatial scale of adaptation (Antonovics and Bradshaw 1970; Galloway and Fenster 2000; Sexton et al. 2013; Anderson et al. 2015; Wright et al. 2017). We were interested in quantifying adaptation across a broad climatic gradient to examine adaptive responses to historical climate and inform predictions of responses to climate change. Across the gradient sampled in our experiment, adaptation in ragweed occurs at a spatial scale of hundreds of kilometers. In IL, the garden with the steepest relationship between fitness and distance, our model predicted fitness to decrease by about one fruit per km; thus, it would take ∼800 km for fitness to decline 50% from its predicted maximum (Fig. 3b). Adaptation at such a large spatial scale likely reflects both the high rates of gene flow among native populations of ragweed (Hämälä et al. 2020), which tend to produce large amounts of wind-dispersed pollen, and the fact that the main agent of selection captured by our sampling design, climate, varies at relatively large scales.

Although the spatial scale of adaptation in ragweed is large, our results indicate that the sheer magnitude of climate change will nonetheless lead to local maladaptation of many ragweed populations. For example, the temperature in our gardens was approximately one degree warmer than historical averages, which is roughly equivalent to moving 165 km south along the latitudinal gradient from which we sampled. In the two most southern gardens, IL and MO, this temperature increase is what presumably led to source populations from warmer climes outperforming local populations. This mirrors the expectation shown in Figure 1b, where peak fitness is shifted away from zero (local populations) and toward populations sourced from warmer climes (note that in our experiment we captured a linear section of the quadratic relationship shown in Figure 1b; with more populations sourced from further south a quadratic relationship may have emerged). Similar signatures of climate change were revealed in a recent meta-analysis of transplant experiments, where population home-site advantage decays with increasing differences between contemporary and historical temperatures (Bontrager et al. 2020). In our experiment, the deviation from climatic norms explains, in part, why directional latitudinal distance was a far better predictor of lifetime fitness than absolute latitudinal (or Euclidean) distance. Climate change has and will continue to push ecological systems toward disequilibrium. As the climate warms, the adaptive differences between populations sourced north and south of a garden will be highly consequential, and the use of directional measures of distance will be necessary to understand shifted patterns of adaptation (Fig. 1b).

Our experiment also uncovered substantial among-garden differences in local adaptation and the life history stages underlying adaptive differentiation. In IA, we found no signal of local adaptation or maladaptation to climate, suggesting fitness variation among populations may be due to selective agents varying at finer scales than those captured in our study. Local adaptation was apparent only in the most northern garden, MN, while local maladaptation was evident in the two southern gardens, IL and MO. However, all gardens experienced similar temperature deviations during the experiment – why was local maladaptation only visible in IL and MO? Our analyses of separate life history stages help to answer this question. In MN, IL, and MO, southern source populations had higher fecundity than northern populations. In IL and MO, this resulted in populations sourced from south of the gardens outperforming local populations, presumably because the southernmost source populations had the greatest capacity to increase fecundity during the warm weather of the experiment. In MN, however, plants from southern populations were much less likely to produce fruit than plants from northern populations because southern populations flowered considerably later in the growing season (Fig. S4), which left little time for fertilization and fruit set before fall frost.

The latitudinal gradient in flowering phenology arises presumably due to adaptive differences in phenological cues based on photoperiod (Allard 1945; Thomas and Vince-Prue 1996; Deen et al. 1998; Leiblein-Wild and Tackenberg 2014; Gorton et al. 2019). Thus, adaptation to photoperiod cues led to the maintenance of local adaptation in MN. However, it is important to note that the strength of local adaptation in MN was modest, with distance from the common garden only explaining about 20 percent of the variation in fitness among populations. Even though the phenological advantage of local populations was strong, the advantage was partially offset by the increased fecundity of southern individuals that fruited. The signal of local adaptation in MN may have been especially weak because the year of our experiment had the latest first frost on record for that locale (MN DNR 2017). The extended growing season likely allowed a higher proportion of southern genotypes to successfully fruit than would have with a typical first frost date, thereby weakening the *fitness ∼ distance* relationship in that garden. Thus, longer frost-free periods due to climate change may increasingly weaken local adaptation of northern ragweed populations.

A common prediction is that as climate warms, populations may shift northward and track the climatic envelope to which they are currently adapted, a prediction that has already borne out for some species (Chen et al. 2011; Rumpf et al. 2018). Our results from the two southern gardens suggest that individuals from southern populations that migrate northward will likely do well across much of the range and may outperform local populations. However, at the northern edge of the range, the growing season may lengthen with climate change, but daylength will not, and the fitness of southern populations migrating northwards will largely depend on adaptive evolution of flowering cues (e.g., Griffith and Watson 2006). Taken together, our results suggest that in response to a changing climate, more southerly populations of ragweed might shift northward, or gene flow from southern populations may accelerate adaptation of more central populations to warmer conditions, potentially resulting in greater fecundity and larger populations. By contrast, without adaptation in phenological cues, daylength at the edge of the range may act as a barrier to range expansion (assuming daylength remains an accurate cue for probability of frost). Thus, climate change could conceivably lead to a contraction of ragweed’s geographic range, as populations migrate northward, but migration at the northern edge is prevented or slowed by constraints on adaptation to photoperiod. These results highlight the important observation that the environment, and thus adaptation, is multivariate (Antonovics 1976; Benning and Moeller 2019), and populations’ responses to rising temperature, either via migration or adaptation, will interact with other important axes of adaptation and will likely differ from predictions based solely on climatic change.

In summary, our work underscores that many populations are not in equilibrium with their environment. Climate change is already disrupting local adaptation, though population responses and the traits and life history stages underlying local adaptation will likely differ across regions of a species’ range. Thus, employing multiple common gardens across a large portion of a species’ range is essential in experiments seeking to understand spatial patterns of adaptation. The inclusion of many source populations is also valuable in such studies, as it enables one to treat distance as a continuous, as opposed to categorical (e.g., home vs. away) predictor in analyses, resulting in a quantifiable estimate of the spatial scale of adaptation as the change in fitness per unit distance. Furthermore, the use of directional, as opposed to absolute measures of distance will be key to identify disrupted patterns of local adaptation. Moving forward, information gleaned from common garden studies like ours can inform better predictions of species range shifts and the responses of populations to climate change.

## Acknowledgements

We thank M. Merello, L. Peters, the Missouri Botanical Garden, R. Noyes, C. Rentschler and J. Carlson for assistance with seed collections. Y.W Lee, S. DelSerra, and J. Yoon Kim assisted with data collection in the field. We also thank the following individuals and associated institutions for use of their field stations: M. Lotsetter at the Rosemount Outreach and Research Center (University of Minnesota), E. Hill at the Conrad Environmental Research Area (Grinnell College), M. Bernards at the University of Western Illinois, and K. Medley at the Tyson Research Center (Washington University). AJG was funded by a Carol H and Wayne Pletcher Fellowship and a Dayton Fellowship (Bell Natural History Museum).

## Literature Cited

Aitken, S. N., S. Yeaman, J. A. Holliday, T. Wang, and S. Curtis-McLane. 2008. Adaptation, migration or extirpation: climate change outcomes for tree populations. Evolutionary Applications 1:95–111.

Allard, H. A. 1945. Flowering Behavior and Natural Distribution of the Eastern Ragweeds (Ambrosia) as Affected by Length of Day. Ecology 26:387–394. Ecological Society of America.

Anderson, J. T. 2015. Plant fitness in a rapidly changing world. New Phytologist, doi: 10.1111/nph.13693.

Anderson, J. T., N. Perera, B. Chowdhury, and T. Mitchell-Olds. 2015. Microgeographic Patterns of Genetic Divergence and Adaptation across Environmental Gradients in Boechera stricta (Brassicaceae)*. The American Naturalist S000–S000.

Anderson, J. T., and S. M. Wadgymar. 2019. Climate change disrupts local adaptation and favours upslope migration. Ecol. Lett. 348.

Antonovics, J. 1976. The Nature of Limits to Natural Selection. Ann. Mo. Bot. Gard. 63:224–247. Missouri Botanical Garden Press.

Antonovics, J., and A. D. Bradshaw. 1970. Evolution in closely adjacent plant populations. VIII. Clinal patterns at a mine boundary. Heredity 25:349–362. cabdirect.org.

Becker, U., G. Colling, P. Dostal, A. Jakobsson, and D. Matthies. 2006. Local adaptation in the monocarpic perennial Carlina vulgaris at different spatial scales across Europe. Oecologia 150:506–518.

Benning, J. W., and D. A. Moeller. 2019. Maladaptation beyond a geographic range limit driven by antagonistic and mutualistic biotic interactions across an abiotic gradient. Evolution 73:2044–2059.

Benning, J. W., and D. A. Moeller. 2021. Microbes, mutualism, and range margins: testing the fitness consequences of soil microbial communities across and beyond a native plant’s range. New Phytol. 229:2886–2900.

Bontrager, M., and A. L. Angert. 2019. Gene flow improves fitness at a range edge under climate change. Evol Lett 3:55–68.

Bontrager, M., C. D. Muir, C. Mahony, D. E. Gamble, R. M. Germain, A. L. Hargreaves, E. J. Kleynhans, K. A. Thompson, and A. L. Angert. 2020. Climate warming weakens local adaptation.

Breheny, P., and W. Burchett. 2017. visreg: Visualization of Regression Models.

Campbell-Staton, S. C., A. Bare, J. B. Losos, S. V. Edwards, and Z. A. Cheviron. 2018. Physiological and regulatory underpinnings of geographic variation in reptilian cold tolerance across a latitudinal cline. Mol. Ecol. 27:2243–2255.

Chauvel, B., F. Dessaint, C. Cardinal-Legrand, and F. Bretagnolle. 2006. The historical spread of Ambrosia artemisiifolia L. in France from herbarium records. Journal of Biogeography 33:665–673.

Chen, I.-C., J. K. Hill, R. Ohlemüller, D. B. Roy, and C. D. Thomas. 2011. Rapid range shifts of species associated with high levels of climate warming. Science 333:1024–1026.

Chevin, L.-M., M. E. Visser, and J. Tufto. 2015. Estimating the variation, autocorrelation, and environmental sensitivity of phenotypic selection. Evolution 69:2319–2332.

Colautti, R. I., and S. C. H. Barrett. 2013. Rapid adaptation to climate facilitates range expansion of an invasive plant. Science 342:364–366. science.sciencemag.org.

de Villemereuil, P., A. Charmantier, D. Arlt, P. Bize, P. Brekke, L. Brouwer, A. Cockburn, S. D. Côté, F. S. Dobson, S. R. Evans, M. Festa-Bianchet, M. Gamelon, S. Hamel, J. Hegelbach, K. Jerstad, B. Kempenaers, L. E. B. Kruuk, J. Kumpula, T. Kvalnes, A. G. McAdam, S. E. McFarlane, M. B. Morrissey, T. Pärt, J. M. Pemberton, A. Qvarnström, O. W. Røstad, J. Schroeder, J. C. Senar, B. C. Sheldon, M. van de Pol, M. E. Visser, N. T. Wheelwright, J. Tufto, and L.-M. Chevin. 2020. Fluctuating optimum and temporally variable selection on breeding date in birds and mammals. Proc. Natl. Acad. Sci. U. S. A. 117:31969–31978.

Deen, W., T. Hunt, and C. J. Swanton. 1998. Influence of temperature, photoperiod, and irradiance on the phenological development of common ragweed (Ambrosia artemisiifolia). Weed Sci. 46:555–560. Cambridge University Press.

Essl, F., K. Biró, D. Brandes, O. Broennimann, J. M. Bullock, D. S. Chapman, B. Chauvel, S. Dullinger, B. Fumanal, A. Guisan, G. Karrer, G. Kazinczi, C. Kueffer, B. Laitung, C. Lavoie, M. Leitner, T. Mang, D. Moser, H. Müller-Schärer, B. Petitpierre, R. Richter, U. Schaffner, M. Smith, U. Starfinger, R. Vautard, G. Vogl, M. von der Lippe, and S. Follak. 2015. Biological Flora of the British Isles: Ambrosia artemisiifolia. Journal of Ecology 103:1069–1098.

Etterson, J. R., and R. G. Shaw. 2001. Constraint to adaptive evolution in response to global warming. Science 294:151–154. science.sciencemag.org.

Fournier-Level, A., A. Korte, M. D. Cooper, M. Nordborg, J. Schmitt, and A. M. Wilczek. 2011. A map of local adaptation in Arabidopsis thaliana. Science 334:86–89.

Fox, J., and S. Weisberg. 2019. An R Companion to Applied Regression. Sage, Thousand Oaks CA.

Galloway, L. F., and C. B. Fenster. 2000. Population differentiation in an annual legume: local adaptation. Evolution 54:1173–1181. Wiley Online Library.

Genton, B. J., J. A. Shykoff, and T. Giraud. 2005. High genetic diversity in French invasive populations of common ragweed, Ambrosia artemisiifolia, as a result of multiple sources of introduction. Molecular Ecology 14:4275–4285.

Gorton, A. J., P. Tiffin, and D. A. Moeller. 2019. Does adaptation to historical climate shape plant responses to future rainfall patterns? A rainfall manipulation experiment with common ragweed. Oecologia 190:941–953.

Griffith, T.M. and Watson, M.A. 2006. Is evolution necessary for range expansion? Manipulating reproductive timing of a weedy annual transplanted beyond its range. The American Naturalist, 167(2), pp.153–164.

Hämälä, T., A. J. Gorton, D. A. Moeller, and P. Tiffin. 2020. Pleiotropy facilitates local adaptation to distant optima in common ragweed (Ambrosia artemisiifolia). PLoS Genet. 16:e1008707. Public Library of Science.

Hargreaves, A. L., R. M. Germain, M. Bontrager, J. Persi, and A. L. Angert. 2020. Local Adaptation to Biotic Interactions: A Meta-analysis across Latitudes. Am. Nat. 195:395–411.

Hereford, J. 2009. A quantitative survey of local adaptation and fitness trade-offs. Am. Nat. 173:579–588. journals.uchicago.edu.

Hodgins, K. A., and L. Rieseberg. 2011. Genetic differentiation in life-history traits of introduced and native common ragweed (Ambrosia artemisiifolia) populations. Journal of Evolutionary Biology 24:2731–2749.

Janson, K. 1983. Selection and migration in two distinct phenotypes of Littorina saxatilis in Sweden. Oecologia 59:58–61.

Jones, K. L. 1936. Studies on Ambrosia I. The inheritance of floral types in the ragweed, Ambrosia elatior L. American Midland Naturalist 17:673–699.

Joshi, J., B. Schmid, M. C. Caldeira, P. G. Dimitrakopoulos, J. Good, R. Harris, A. Hector, K. Huss-Danell, A. Jumpponen, A. Minns, C. P. H. Mulder, J. S. Pereira, A. Prinz, M. Scherer-Lorenzen, a. S. D. Siamantziouras, a. C. Terry, a. Y. Troumbis, and J. H. Lawton. 2001. Local adaptation enhances performance of common plant species. Ecology Letters 4:536–544.

Kartesz, J. T. 2013. The Biota of North America Program (BONAP).

Kawecki, T. J., and D. Ebert. 2004. Conceptual issues in local adaptation. Ecol. Lett. 7:1225–1241. Blackwell Science Ltd.

Keller, S. R., N. Levsen, P. K. Ingvarsson, M. S. Olson, and P. Tiffin. 2011. Local selection across a latitudinal gradient shapes nucleotide diversity in balsam poplar, Populus balsamifera L. Genetics 188:941–952.

Kostanecki, A., A. J. Gorton, and D. A. Moeller. 2021. An urban-rural spotlight: Evolution at small spatial scales among urban and rural populations of common ragweed. Journal of Urban Ecology 7:1–8.

Lacey, E. P. 1988. Latitudinal variation in reproductive timing of a short-lived monocarp, Daucus carota (Apiaceae). Ecology 69:220–232.

Leiblein-Wild, M. C., and O. Tackenberg. 2014. Phenotypic variation of 38 European Ambrosia artemisiifolia populations measured in a common garden experiment. Biological Invasions 16:2003–2015.

Lenoir, J., R. Bertrand, L. Comte, L. Bourgeaud, T. Hattab, J. Murienne, and G. Grenouillet. 2020. Species better track climate warming in the oceans than on land. Nat Ecol Evol, doi: 10.1038/s41559-020-1198-2.

Lenormand, T. 2002. Gene flow and the limits to natural selection. Trends Ecol. Evol. 17:183–189.

Lenth, R. 2018. Emmeans: Estimated marginal means, aka least-squares means. R Package Version 1.

Li, C., O. Junttila, A. Ernstsen, P. Heino, and E. T. Palva. 2003. Photoperiodic control of growth, cold acclimation and dormancy development in silver birch (Betula pendula) ecotypes. Physiologia Plantarum 117:206–212.

Linhart, Y. B., and M. C. Grant. 1996. Evolutionary Significance of Local Genetic Differentiation in Plants. Annual Review of Ecology and Systematics 27:237–277.

MN DNR. 2017. https://www.dnr.state.mn.us/climate/journal/1610_frost.html.

Montalvo, A. M., and N. C. Ellstrand. 2000. Transplantation of the subshrub Lotus scoparious: testing the home-site advantage hypothesis. Conservation Biology 14:1034–1045.

O’Brien, E. K., M. Higgie, A. Reynolds, A. A. Hoffmann, and J. R. Bridle. 2017. Testing for local adaptation and evolutionary potential along altitudinal gradients in rainforest Drosophila: beyond laboratory estimates. Glob. Chang. Biol. 23:1847–1860. Oregon State University. 2020. PRISM Climate Group.

R Foundation For Statistical Computing. 2021. R: A language and environmnet for statistical computing.

Richardson, J. L., M. C. Urban, D. I. Bolnick, and D. K. Skelly. 2014. Microgeographic adaptation and the spatial scale of evolution. Trends Ecol. Evol. 29:165–176.

Rumpf, S. B., K. Hülber, G. Klonner, D. Moser, M. Schütz, J. Wessely, W. Willner, N. E. Zimmermann, and S. Dullinger. 2018. Range dynamics of mountain plants decrease with elevation. Proc. Natl. Acad. Sci. U. S. A. 115:1848–1853.

Runquist, R. D. B., A. J. Gorton, J. B. Yoder, N. J. Deacon, J. J. Grossman, S. Kothari, M. P. Lyons, S. N. Sheth, P. Tiffin, and D. A. Moeller. 2020. Context dependence of local adaptation to abiotic and biotic environments: A quantitative and qualitative synthesis. American Naturalist 195:412–431.

Sambatti, J. B. M., and K. J. Rice. 2006. Local adaptation, patterns of selection, and gene flow in the Californian serpentine sunflower (Helianthus exilis). Evolution; international journal of organic evolution 60:696–710.

Savolainen, O., T. Pyhäjärvi, and T. Knürr. 2007. Gene Flow and Local Adaptation in Trees. Annual Review of Ecology, Evolution, and Systematics 38:595–619.

Sexton, J. P., S. B. Hangartner, and A. A. Hoffmann. 2013. Genetic Isolation By Environment or Distance: Which Pattern of Gene Flow Is Most Common? Evolution 68:1–15.

Slatkin, M. 1987. Gene flow and the geographic structure of natural populations. Science 236:787–792.

Slatkin, M. 1985. Gene flow in natural populations. Annu. Rev. Ecol. Syst. 16:393–430. Annual Reviews 4139 El Camino Way, PO Box 10139, Palo Alto, CA 94303-0139, USA.

Thomas, B., and D. Vince-Prue. 1996. Photoperiodism in Plants. Elsevier.

Toole, E. H., and E. Brown. 1946. Final results of the Duvel buried seed experiment. Journal of Agricultural Research 72:201–210.

Whitlock, M. C. 2015. Modern Approaches to Local Adaptation. Am. Nat. 186 Suppl 1:S1–4.

Wickham, H., M. Averick, J. Bryan, W. Chang, L. McGowan, R. François, G. Grolemund, A. Hayes, L. Henry, J. Hester, M. Kuhn, T. Pedersen, E. Miller, S. Bache, K. Müller, J. Ooms, D. Robinson, D. Seidel, V. Spinu, K. Takahashi, D. Vaughan, C. Wilke, K. Woo, and H. Yutani. 2019. Welcome to the tidyverse. J. Open Source Softw. 4:1686. The Open Journal.

Wilczek, A. M., M. D. Cooper, T. M. Korves, and J. Schmitt. 2014. Lagging adaptation to warming climate in Arabidopsis thaliana. Proc. Natl. Acad. Sci. U. S. A. 111:7906–7913.

Williams, G. C. 1966. Adaptation and natural selection. Princeton University Press, Princeton, NJ, USA.

Wright, S. J., D. Cui Zhou, A. Kuhle, and K. M. Olsen. 2017. Continent-Wide Climatic Variation Drives Local Adaptation in North American White Clover. J. Hered. 109:78–89.

Xu, H., S. Qiang, Z. Han, J. Guo, Z. Huang, H. Sun, S. He, H. Ding, H. Wu, and F. Wan. 2006. The status and causes of alien species invasion in China. Biodiversity and Conservation 15:2893–2904.

