## Supplementary material for "The spatial scale of adaptation in a native annual plant and its implications for responses to climate change": SI

***Predicting fitness***

Measuring female fitness on all plants was not possible in the field due to logistical constraints. However, we did collect extensive phenotypic data preceding fruit set, from planting until 16 weeks post planting, for all plants. Thus, we used the full lifetime fitness data we had from the subset of 9 populations to build models based on individual plant phenotypes that predicted the two final fitness components of interest: probability of producing fruit, and number of fruits. We used these models to predict P(fruiting) and number of fruits for the remaining plants in the data set (those from the other 13 populations, and individuals from the 9 populations whose fitness was not estimated in the field). Final data set sizes were N = 524 for the model building data set (hereafter, modeling set) and N = 1130 for the prediction data set (hereafter, prediction set). These datasets excluded any plants that we know died (N = 530), because their fitness was known to be zero. We chose models that maximized the amount of variance in fitness components that was explained, while minimizing the prediction of outliers in the predicted data.

We used logistic regression to model P(fruiting) as predicted by garden, plant developmental stage at 12 weeks, and their interaction, with model formulation *glm(fruiting ~ Garden X Stage_12wk, family = binomial)*. This model correctly predicted fruiting in 508/524 individuals (97%), after rounding estimated probabilities to 0 or 1. We then used this model to predict P(fruiting) for all individuals in the prediction set, rounding this probability to 0 or 1.

We used linear regression to model fruit number as predicted by garden, height at 16 weeks, developmental stage at 12 weeks, and their interactions, and the interaction of garden and female flowering stage at 12 weeks; the model took the form *lm(fruits ~ Garden X Height_16wk X Stage_12wk + Garden:Female_12wk)*. Model $R^2 = 0.38$. Though the model had a modest $R^2$, what we ultimately were interested in was whether these individual level phenotypic models could accurately predict mean population values, as this was the unit of analysis in our main analyses. The correlation between predicted (based on the *fruits* model above) and observed population mean number of fruits in the modeling set of 9 focal populations was high, $r = 0.94$. Examining the relationship between one of our distance metrics, directional latitudinal distance, and the observed vs. predicted mean population fruit set, we can see the data sets produce extremely similar regression results (Fig. S1). The correlation between predicted population mean fitness (based on individual fitness as predicted by the product of P(fruiting) and number of fruits) and observed population mean fitness in the modeling set of 9 focal populations was also high, $r = 0.94$. We used this *fruits* model to predict the number of fruits produced for all individuals in the prediction set.

After predicting P(fruting) and number of fruits for all individuals in the prediction set, we calculated fitness for those individuals as the product of those two numbers. These predictions produced one outlier at MN, whose predicted fitness, due to the plant's extremely large size, was almost four times that of most fecund individual measured in that garden (though similar to some individuals measured at other gardens). We kept this outlier in the data set because it had no qualitative effect on our analyses (regression of *fitness ~ directional latitudinal distance* at MN garden: with outlier, slope = 0.135 fruits per km, $P_{1,20} = 0.027$, $R^2 = 0.18$; without outlier, slope = 0.10 fruits per km, $P_{1,20} = 0.038$, $R^2 = 0.16$.)

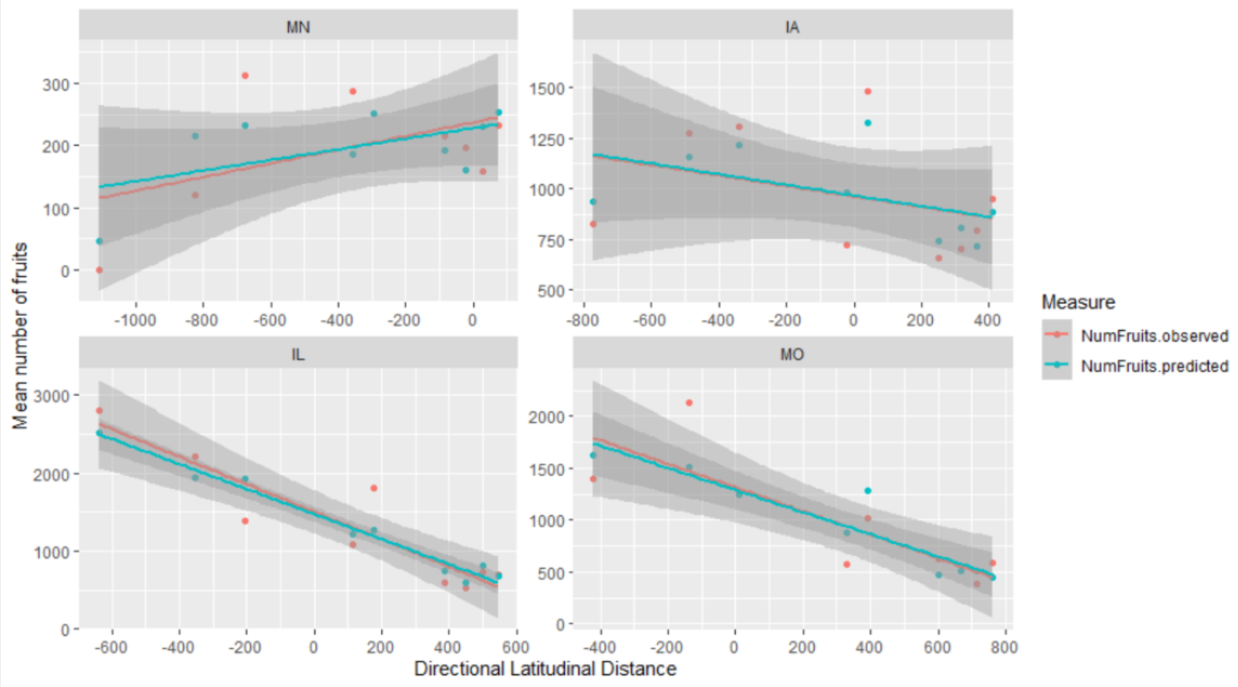


**Figure S1.** Population mean number of fruits regressed on directional latitudinal distance at each garden, for surviving individuals in the nine “focal” populations. Red points and lines are observed values and the associated linear regression; teal points and lines are predicted values and linear regression.

Of course, the important assumption in this approach is that the modeled relationship between earlier phenotypic data and fruit set in the 9 “focal” populations for which we had fruit set data holds for individuals from the 13 “extra” populations for which we did not record final fitness data. As a proof of concept, we took a similar approach to the one we used for fruit set to model the probability of a plant having any mature fruits at 16 weeks, which was phenotypic data we recorded for every surviving plant in the field and the latest life history data we had for every plant. We modeled the probability of a plant having any mature fruits at 16 weeks as predicted by garden, female flowering stage at 12 weeks, height at 12 weeks, and their interactions, with logistic regression, with model formulation *glm(fruiting_16wk ~ Garden X*

*FemaleStage_12wk X Height_12wk, family = binomial)*. We built this model using individuals from the nine focal populations, and then used it to predict the probability of fruiting at 16 weeks for individuals in the 13 extra populations, rounding probabilities to 0 or 1. These predictions matched the observed data in the 13 extra populations very well (Fig. S2). Overall in the 13 extra populations, we observed 18.2% of surviving plants had mature fruits at 16 weeks; model predictions resulted in an estimate of 16%. A two-proportions Z-test between the predicted and observed data indicate no significant difference ($P$ = 0.22)

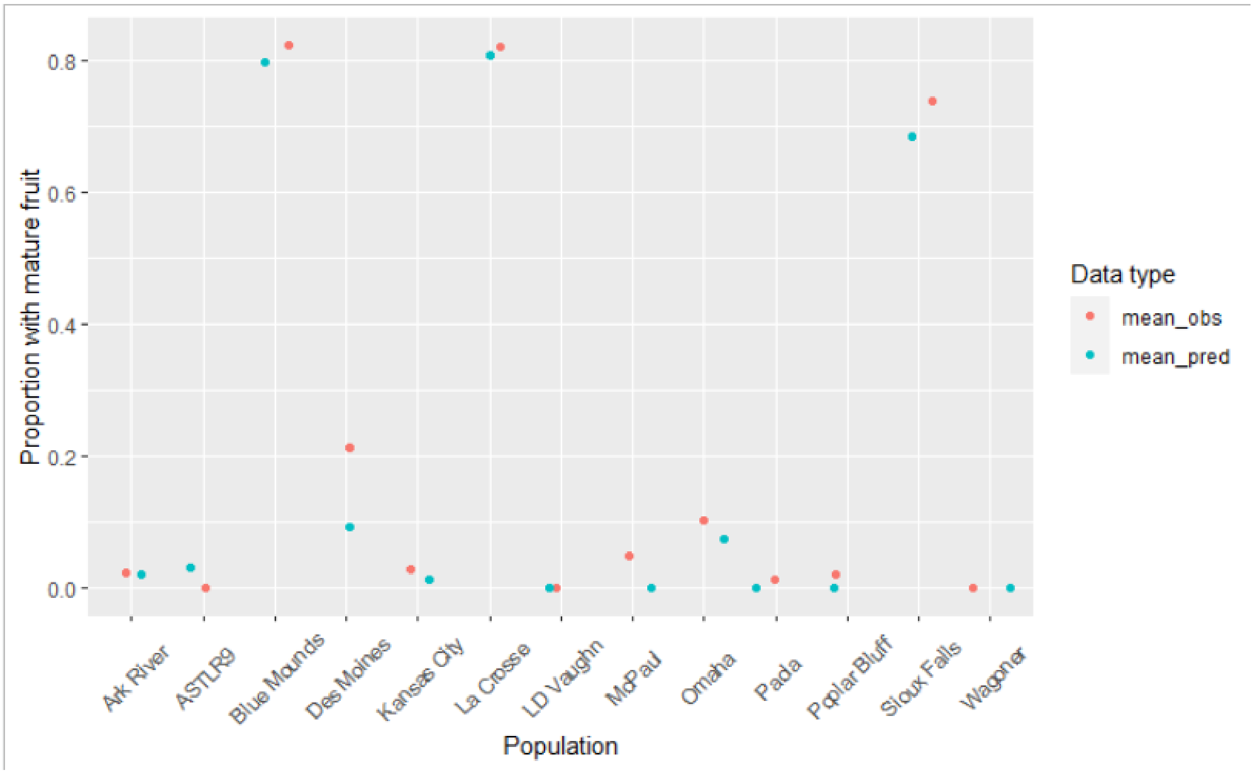


**Figure S2.** Proportion of surviving plants with mature fruit at 16 weeks for the 13 "extra" populations, based on observed data (red points) and values predicted from a model built using the 9 "focal" populations (teal points).

We can also ask whether the relationship between fitness and distance seems to change between the two sets of populations, or whether using the full data set with predictions changes our inference relative to the data set with only the nine populations for which fruit set was measured in the field. In Figure S3, we can see that regardless of the data set used (9 “focal” populations, 13 “extra” populations, or all 22 populations combined), the relationship between directional latitudinal distance and fitness remains similar. To explicitly test whether the relationship between fitness and distance differed using “focal” vs. “extra” data sets, we built regression models of *fitness* ~ *distance* * *data set* for Euclidean, absolute latitudinal, directional latitudinal, absolute temperature, and directional temperature distance at each garden. A significant *distance* × *data set* interaction would indicate that the relationship between fitness and distance varied between data sets. None of the distance metrics showed significant interactions with data set at any garden (all $P > 0.2$; see archived R code for full ANOVA tables).

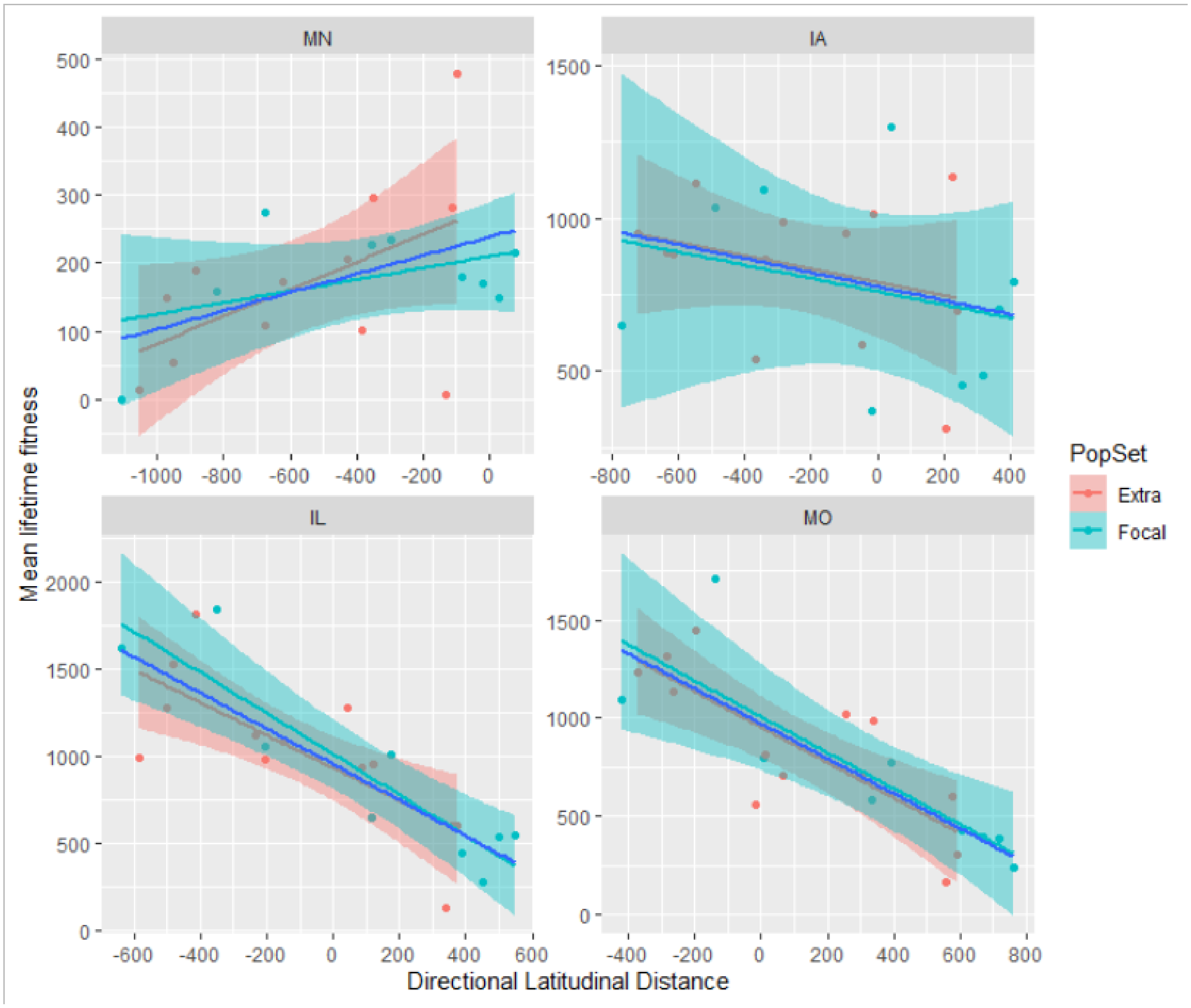

**Figure S3.** Mean lifetime fitness regressed on directional latitudinal distance. Teal points and linear regression lines/bands are based on the “focal” 9 populations only; red points and linear regression lines/bands are based on the “extra” 13 populations only; the blue linear regression (no band) is based on the full data set with 22 populations.

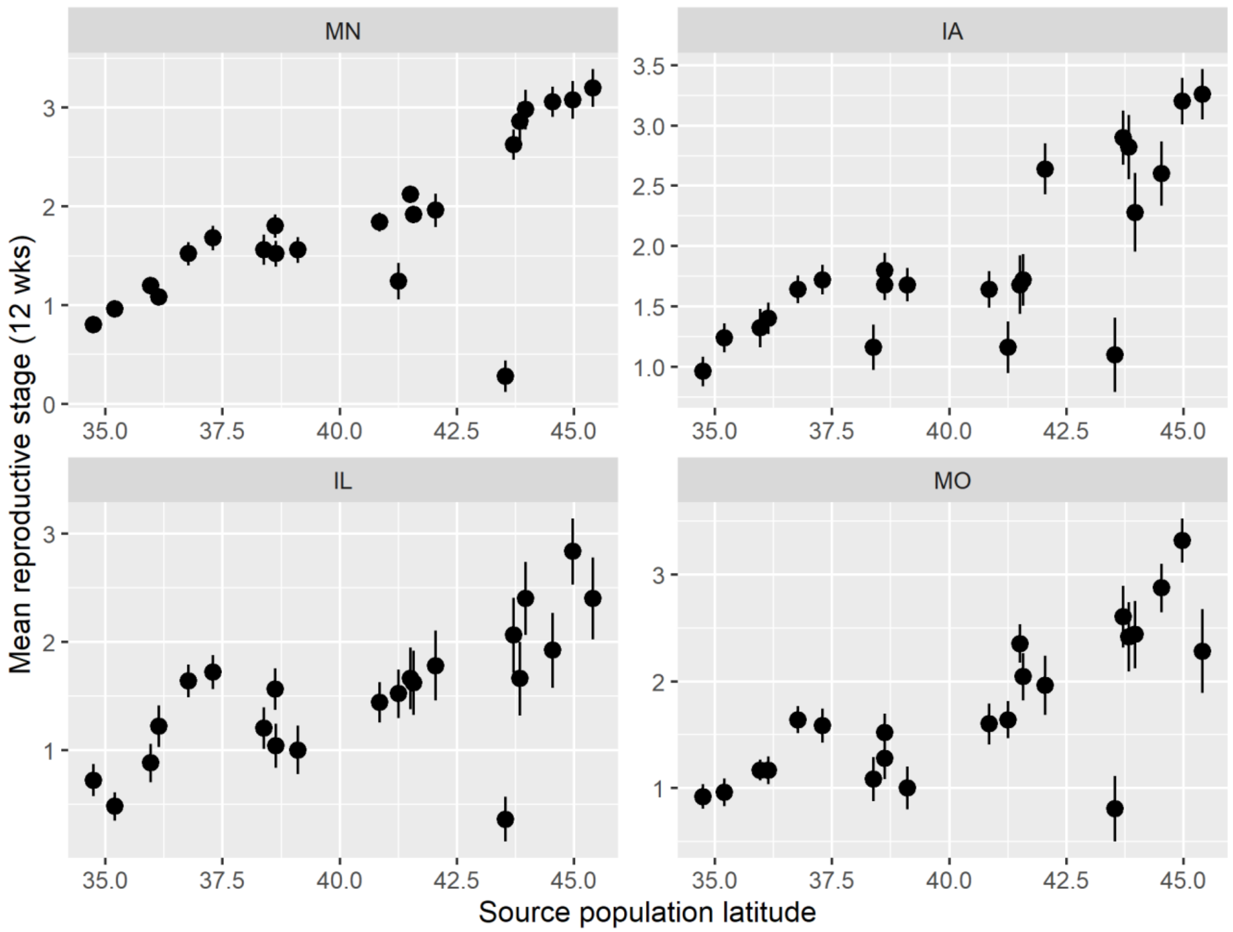


**Figure S4**. The effect of source population latitude on developmental stage at 12 weeks (mean ± SE) at each common garden. Values above 3 indicate either male or female flowers were open. See Table S3 for a description of all potential stages.

### SI Tables

**Table S1**. Description of populations included in the latitudinal reciprocal transplant experiment. Populations are listed in ascending order by latitude. Temperature and precipitation indicate population growing season mean temperature and mean cumulative precipitation (averaged 1986-2015). *n* indicates the number of maternal plants that were included from that population. Populations that had a fruit count in the field are indicated with a "Y".

| Population | Latitude | Longitude | Temperature (C) | Precipitation (mm) | *n* | Fruit counts |
|---|---|---|---|---|---|---|
| LTR | 34.74136566 | -92.25947634 | 24.05 | 95.43 | 23 | Y |
| LDV | 35.19796198 | -91.64370698 | 23.54 | 93.65 | 25 | |
| WAG | 35.96535395 | -95.45782392 | 23.31 | 105.57 | 25 | |
| ARK | 36.13697554 | -95.99441688 | 23.48 | 103.62 | 19 | |
| POP | 36.76623098 | -90.40547492 | 22.31 | 98.55 | 15 | |
| CLW | 37.29567794 | -90.36880106 | 20.54 | 99.81 | 25 | Y |
| SR9 | 38.38645 | -89.92686 | 21.36 | 94.11 | 25 | |
| SU8 | 38.61736 | -90.26415 | 22.01 | 97.98 | 25 | Y |
| PAO | 38.62834563 | -94.81757954 | 21.21 | 113.47 | 25 | |
| KAC | 39.10268604 | -94.60068525 | 21.63 | 116.49 | 25 | |
| MPL | 40.84571607 | -95.80212162 | 19.74 | 99.18 | 25 | |
| OMA | 41.25688838 | -95.94715796 | 19.72 | 95.26 | 12 | |
| ROI | 41.51026297 | -90.56311849 | 19.75 | 100.15 | 23 | Y |
| DMS | 41.57788835 | -93.62835764 | 19.57 | 97.22 | 25 | |
| PRC | 42.05130841 | -90.63232642 | 17.60 | 95.07 | 25 | Y |
| SFA | 43.53882148 | -96.72653811 | 17.54 | 79.38 | 24 | |
| BMO | 43.71449138 | -96.18077593 | 17.06 | 86.49 | 25 | |
| LAC | 43.83766655 | -91.2513499 | 18.34 | 95.63 | 25 | Y |
| LEW | 43.96234593 | -91.83442563 | 16.38 | 98.62 | 18 | |
| R10 | 44.53656 | -93.02021 | 17.19 | 99.41 | 25 | Y |
| U25 | 44.96804 | -93.22023 | 17.93 | 96.40 | 18 | Y |
| R16 | 45.40264 | -93.19718 | 16.57 | 94.71 | 25 | Y |

**Table S2.** Location of common gardens and garden growing season temperature and precipitation during the experiment year (2016).

| Common garden | Latitude | Longitude | Temperature (C) | Precipitation (mm) | Hosting Institution |
|---|---|---|---|---|---|
| **MN** | 44.71645 | -93.09789 | 18.48 | 120.74 | Rosemount Outreach and Research Center, University of Minnesota |
| **IA** | 41.68763 | -92.86527 | 19.24 | 105.74 | Conrad Environmental Research Area, Grinnell |
| **IL** | 40.47617 | -90.6844 | 20.43 | 105.60 | Western Illinois Agricultural Station, Western Illinoi University |
| **MO** | 38.53701 | -90.5589 | 22.84 | 126.39 | Tyson Research Center, Washington University |

**Table S3.** Vegetative and reproductive scoring that was used on each individual plant at weeks 4,8,12 and 16. Note that an individual could receive an intermediate score between stages.

| Stage | Developmental stage | Male flowering | Female flowering |
|---|---|---|---|
| 1 | Vegetative; only leaves | Male inflorescences developing, only buds visible | Stigmas visible, no immature fruit |
| 2 | Initiation of reproduction | Male inflorescences visible, but flowers closed | Stigmas visible, ~<25% are immature fruit |
| 3 | Male or female flowers open | <50% flowers open per inflorescence | <50% are immature fruit, few stigmas |
| 4 | Fruit visible | >50% flowers open per inflorescence | >50% immature fruit, ~25% mature fruit |
| 5 | Senesced | All male flowers open or flowers senesced | All mature fruit or senesced |

**Table S4.** Pearson correlation (*r*) among distance metrics used in the analyses, for all gardens together (top row in each cell) and individual gardens.

| | Euclidean | Absolute Latitudinal | Directional Latitudinal | Absolute Longitudinal | Directional Longitudinal | Directional Temperature | Absolute Temperature | Directional Precipitation |
|---|---|---|---|---|---|---|---|---|
| **Absolute Latitudinal** | All: 0.93<br>MN: 0.99<br>IA: 0.96<br>IL: 0.72<br>MO: 0.86 | | | | | | | |
| **Directional Latitudinal** | All: -0.37<br>MN: -0.99<br>IA: -0.74<br>IL: -0.13<br>MO: 0.64 | All: -0.41<br>MN: -1<br>IA: -0.67<br>IL: -0.18<br>MO: 0.7 | | | | | | |
| **Absolute Longitudinal** | All: 0.36<br>MN: 0.36<br>IA: 0.05<br>IL: 0.58<br>MO: 0.57 | All: 0.02<br>MN: 0.26<br>IA: -0.18<br>IL: -0.1<br>MO: 0.1 | All: 0.07<br>MN: -0.3<br>IA: -0.24<br>IL: 0.1<br>MO: 0.11 | | | | | |
| **Directional Longitudinal** | All: -0.15<br>MN: 0.1<br>IA: 0.03<br>IL: -0.61<br>MO: -0.63 | All: 0.11<br>MN: 0.14<br>IA: 0.11<br>IL: 0.06<br>MO: -0.16 | All: -0.33<br>MN: -0.14<br>IA: -0.14<br>IL: -0.14<br>MO: -0.14 | All: -0.68<br>MN: -0.1<br>IA: -0.29<br>IL: -0.99<br>MO: -0.99 | | | | |
| **Directional Temperature** | All: 0.32<br>MN: 0.95<br>IA: 0.71<br>IL: 0.15<br>MO: -0.59 | All: 0.35<br>MN: 0.96<br>IA: 0.65<br>IL: 0.16<br>MO: -0.68 | All: -0.95<br>MN: -0.96<br>IA: -0.96<br>IL: -0.96<br>MO: -0.96 | All: -0.06<br>MN: 0.22<br>IA: 0.18<br>IL: -0.04<br>MO: -0.05 | All: 0.29<br>MN: 0.08<br>IA: 0.08<br>IL: 0.08<br>MO: 0.08 | | | |
| **Absolute Temperature** | All: 0.72<br>MN: 0.87<br>IA: 0.86<br>IL: 0.6<br>MO: 0.7 | All: 0.76<br>MN: 0.88<br>IA: 0.88<br>IL: 0.75<br>MO: 0.8 | All: 0.07<br>MN: -0.87<br>IA: -0.6<br>IL: 0.2<br>MO: 0.93 | All: 0.06<br>MN: 0.06<br>IA: -0.12<br>IL: 0.06<br>MO: 0.08 | All: -0.09<br>MN: 0.05<br>IA: 0.02<br>IL: -0.12<br>MO: -0.12 | All: -0.13<br>MN: 0.86<br>IA: 0.57<br>IL: -0.23<br>MO: -0.97 | | |
| **Directional Precipitation** | All: -0.1<br>MN: 0.29<br>IA: 0.04<br>IL: -0.17<br>MO: -0.32 | All: -0.1<br>MN: 0.36<br>IA: 0.14<br>IL: -0.19<br>MO: -0.52 | All: -0.23<br>MN: -0.36<br>IA: -0.36<br>IL: -0.36<br>MO: -0.36 | All: -0.08<br>MN: -0.23<br>IA: -0.2<br>IL: -0.01<br>MO: -0.01 | All: 0.05<br>MN: 0<br>IA: 0<br>IL: 0<br>MO: 0 | All: 0.41<br>MN: 0.41<br>IA: 0.41<br>IL: 0.41<br>MO: 0.41 | All: -0.23<br>MN: 0.31<br>IA: 0.11<br>IL: -0.35<br>MO: -0.48 | |
| **Absolute Precipitation** | All: 0.08<br>MN: -0.29<br>IA: -0.16<br>IL: 0.14<br>MO: 0.32 | All: 0.07<br>MN: -0.36<br>IA: -0.23<br>IL: -0.02<br>MO: 0.52 | All: 0.23<br>MN: 0.36<br>IA: 0.37<br>IL: 0.37<br>MO: 0.36 | All: 0.12<br>MN: 0.23<br>IA: 0.24<br>IL: 0.25<br>MO: 0.01 | All: -0.11<br>MN: 0<br>IA: -0.23<br>IL: -0.24<br>MO: 0 | All: -0.41<br>MN: -0.41<br>IA: -0.4<br>IL: -0.4<br>MO: -0.41 | All: 0.22<br>MN: -0.31<br>IA: -0.15<br>IL: 0.19<br>MO: 0.48 | All: -0.94<br>MN: -1<br>IA: -0.7<br>IL: -0.68<br>MO: -1 |

**Table S5.** ANOVA tables for linear regression models at each garden, with separate models for Euclidean, absolute geographic (decomposed into latitude and longitude) directional geographic (latitude and longitude), and absolute and directional climatic (temperature and precipitation) distances as predictors.

| | | MN | | IA | | IL | | MO | |
|---|---|---|---|---|---|---|---|---|---|
| | Term | *F* | *P* | *F* | *P* | *F* | *P* | *F* | *P* |
| **Euclidean** | *Distance (km)* | $6.25_{1,20}$ | 0.021 | $0.22_{1,20}$ | 0.647 | $0.00_{1,20}$ | 0.969 | $8.96_{1,20}$ | 0.007 |
| **Absolute geographic** | *Latitude* | $5.76_{1,19}$ | 0.027 | $0.35_{1,19}$ | 0.563 | $0.00_{1,19}$ | 0.952 | $8.66_{1,19}$ | 0.008 |
| | *Longitude* | $0.11_{1,19}$ | 0.745 | $0.33_{1,19}$ | 0.570 | $0.58_{1,19}$ | 0.455 | $0.18_{1,19}$ | 0.677 |
| **Directional geographic** | *Latitude* | $7.77_{1,19}$ | 0.012 | $2.06_{1,19}$ | 0.167 | $41.61_{1,19}$ | <0.001 | $37.97_{1,19}$ | <0.001 |
| | *Longitude* | $3.73_{1,19}$ | 0.069 | $0.03_{1,19}$ | 0.862 | $0.63_{1,19}$ | 0.438 | $0.04_{1,19}$ | 0.842 |
| | *Latitude*$^2$ | $2.86_{1,18}$ | 0.108 | $0.66_{1,18}$ | 0.427 | $0.81_{1,18}$ | 0.380 | $0.25_{1,18}$ | 0.622 |
| | *Longitude*$^2$ | $0.04_{1,18}$ | 0.844 | $0.02_{1,18}$ | 0.884 | $0.08_{1,18}$ | 0.780 | $0.04_{1,18}$ | 0.852 |
| **Absolute climatic** | *Temperature* | $10.03_{1,19}$ | 0.005 | $0.21_{1,19}$ | 0.656 | $1.84_{1,19}$ | 0.191 | $16.21_{1,19}$ | <0.001 |
| | *Precipitation* | $1.24_{1,19}$ | 0.279 | $0.52_{1,19}$ | 0.478 | $5.05_{1,19}$ | 0.037 | $0.02_{1,19}$ | 0.894 |
| **Directional climatic** | *Temperature* | $5.55_{1,19}$ | 0.029 | $0.63_{1,19}$ | 0.438 | $20.12_{1,19}$ | <0.001 | $16.14_{1,19}$ | 0.001 |
| | *Precipitation* | $1.16_{1,19}$ | 0.295 | $0.42_{1,19}$ | 0.526 | $0.24_{1,19}$ | 0.630 | $0.04_{1,19}$ | 0.845 |
| | *Temperature*$^2$ | $2.27_{1,18}$ | 0.149 | $0.08_{1,18}$ | 0.784 | $0.57_{1,18}$ | 0.461 | $0.23_{1,18}$ | 0.634 |
| | *Precipitation*$^2$ | $2.62_{1,18}$ | 0.122 | $0.20_{1,18}$ | 0.657 | $2.91_{1,18}$ | 0.105 | $2.28_{1,18}$ | 0.148 |

**Table S6.** ANOVA tables for life history stage analyses at each common garden. Directional latitudinal distance was the only predictor in each model. Early survival and fruiting were modeled with logistic regression; fruit number was modeled with simple linear regression.

| | ***P*(Early survival)** | | ***P*(Fruiting)** | | **Fruit number** | |
|---|---|---|---|---|---|---|
| | $\chi^2$ | *P* | $\chi^2$ | P | *F* | *P* |
| **MN** | 4.86 | 0.027 | 252.69 | < 0.001 | 5.26 | 0.033 |
| **IA** | 3.82 | 0.051 | 1.07 | 0.302 | 1.42 | 0.247 |
| **IL** | 0.36 | 0.548 | 0.41 | 0.521 | 118.38 | < 0.001 |
| **MO** | 4.31 | 0.038 | 0.03 | 0.870 | 50.70 | < 0.001 |
